# Aerotolerant Capacity of the Lung Commensal *Prevotella melaninogenica*

**DOI:** 10.64898/2026.01.19.700327

**Authors:** Claire Albright, Gouri Anil, Jacob Evans, Souzane Ntamubano, Ariangela Kozik

**Author notes:** Denotes corresponding author (Ariangela Kozik).

## Abstract

*Prevotella melaninogenica* is a core member of the human oral and respiratory microbiomes, often representing more than 10% of microbial populations in both healthy and diseased lungs. Despite its prevalence in these oxygenated environments, *P. melaninogenica* has been historically classified as a strict obligate anaerobe, ostensibly unable to survive oxygen concentrations exceeding 0.05%. This creates a fundamental biological paradox, as the organism consistently persists in the lower respiratory tract where oxygen levels reach higher than what is tolerated by obligate anaerobes. In this study, we resolve this contradiction by evaluating the growth and tolerance of *P. melaninogenica* across intermediate oxygen concentrations of 2%, 5%, and 8%. Contrary to the long-standing classification, we demonstrate that *P. melaninogenica* maintains growth at 2% and 5% oxygen- a level significantly higher than previously reported for the genus- and exhibits robust aerotolerance in 21% O_2_. Transcriptional profiling via RNA-sequencing reveals that this survival is likely driven by the robust expression of oxidative stress defense and DNA repair machinery. Ultimately, these results provide the first evidence of the specialized mechanisms that enable *Prevotella melaninogenica* to adapt to and colonize the respiratory tract, providing a clearer understanding of its persistence in oxygen-exposed human niches.

**IMPORTANCE:** This study provides a correction to the 100-year-old classification of *Prevotella melaninogenica* as a strict obligate anaerobe. We demonstrate that this key member of the human microbiome is capable of robust growth under oxygen levels previously thought to be lethal. By identifying transcriptional responses associated with growth and survival, we predict how *Prevotella melaninogenica* dominates the oxygenated niches of the respiratory tract. This work reveals the putative mechanisms driving the adaptive evolution of *Prevotella melaninogenica* and its role in human airway ecology.

## INTRODUCTION

*Prevotella melaninogenica* is classified as a Gram-negative bacterium, typically appearing as non-motile short rods or coccobacilli^1^. Since its initial isolation by Oliver and Wherry in 1921, this organism has garnered increasing attention, especially with the development of cultivation-independent approaches that have revealed its significant presence within the human respiratory tract microbiome. Estimates suggest that *P. melaninogenica* comprises around 4–8% of the oral microbiome^1^ and is highly represented in the lung microbiome, accounting for roughly 10% of microbial populations in healthy lungs and up to 13% in individuals with respiratory disease^2^. Research has implicated *P. melaninogenica* not only in respiratory health^3^, but also in a spectrum of respiratory disorders, including long-term COVID, gastroesophageal reflux disease^4,5^, chronic obstructive pulmonary disease (COPD)^6^, and cystic fibrosis^7^.

The frequent detection of *P. melaninogenica* in the lungs—a highly oxygenated environment with approximately 9.9% oxygen^8^—contrasts sharply with traditional views that describe this organism as a strict obligate anaerobe, unable to survive in oxygen concentrations exceeding 0.05%. This paradox exposes fundamental gaps in knowledge regarding the organism’s mechanisms for survival and persistence within the lower respiratory tract.

Historically, studies have typically characterized *P. melaninogenica* as strictly anaerobic; however, these conclusions were based on assessments limited to binary comparisons between fully anaerobic and ambient air conditions, without consideration for intermediate oxygen concentrations^9^. Consequently, the bacterium’s potential for growth in intermediate-oxygen environments and its aerotolerance remain unrecognized. Within the same family, *Prevotellaceae*, related species such as *Bacteroides thetaiotaomicron and Bacteroides fragilis* have demonstrated the ability to proliferate under minimal oxygen levels of 0.14% O_2_ ^10,11^. However, B. fragilis has been found to contain an extremely robust aerotolerance, able to tolerate, but not grow, in 21% O_2_ for 48 hours^12,13,14^. Furthermore, B. fragilis has been found to possess several oxidative stress defense mechanisms similar to those found in facultative anaerobes, and is predicted to be equipped with components of the electron transport chain^11,14,15,16^. To date, no studies have evaluated the growth of *P. melaninogenica* within the intermediate oxygen concentrations characteristic of the lower respiratory tract (1%–9.9% O_2_)^8^, coupled with significant aerotolerance to ambient air encountered in the upper respiratory tract. With these considerations in mind, the present investigation aims to address the extent to which *P. melaninogenica* tolerates and grows in oxic environments. Additionally, we seek to elucidate possible oxygen defense strategies employed by this organism, informed by recent findings from related nanaerobic bacteria.

## RESULTS

Our objective was to determine the growth dynamics and aerotolerance profile of *P. melaninogenica* in low-oxygen conditions. To do this, the inoculating culture of *P. melaninogenica* was initially grown in Brain Heart Infusion (BHI) broth at 37°C under strictly anaerobic conditions (0% O_2_). Inoculums derived from this culture were then cultured in BHI broth at 37°C across a range of oxygen conditions: 2%, 5%, 8% and 21% oxygen. For direct comparison of growth rate, cell density (as determined by culture optical density [OD]), and cell viability over time to an anaerobic control, parallel cultures were simultaneously set up and maintained under 0% O_2_. Growth was quantified by measuring optical density over time, and the number of viable cells was determined by anaerobic plating to quantify colony-forming units (CFU). The identity of the bacteria was confirmed using Gram stain and PCR. We first tested the 2% O_2_ condition against the anaerobic control (Fig. 1A). Both cultures had identical inoculums, as indicated by the culture CFU at hour 0. The replication rate of the culture at 0% O_2_ was 1.29 hours, whereas in 2% O_2_ the replication rate was 2.80 hours. Due to the slower replication rate in 2% O_2_, the culture reached its maximum optical density 12 hours later; however, there was no significant difference in the maximum optical density of the two cultures. Furthermore, at the peak optical density, there was no significant difference in the two culture CFU. Next, to assess the impact of a moderate increase in oxygen, we tested 5% O_2_ vs 0% O_2_ (Fig. 1B). The replication rate at 0% O_2_ was 1.20 hours, whereas in 5% O_2_ the replication rate was 1.59 hours. The culture in 5% O_2_ reached its maximum optical density 6 hours later than the culture in 0% O_2_. Strikingly, this is 6 hours faster than when grown at 2% O_2,_ reflecting the quicker replication rate in 5% when compared to the 2% cultures. Similar to the 2% O_2_ results, there was no significant difference in the maximum optical density of 5% O_2_ and 0% O_2_ cultures. We then tested 8% O_2_ vs 0% O_2_ (Fig. 1C). *P. melaninogenica* cultures did not grow in 8% O_2_ over the course of 24 hours, indicating that the upper threshold for growth falls between 5% and 8% O_2_. The culture CFU at hour 0 was ∼10^7^.^3^, confirming initial cell viability. After 24 hours, the culture CFU significantly decreased to ∼10^5^.^5^, but was not reduced to zero. Finally, we evaluated growth and viability under fully aerobic conditions, testing 21% O_2_ vs 0% O_2_ (Fig. 1D). As expected, *P. melaninogenica cultures* did not grow in 21% O_2_ over the course of 24 hours. The culture CFU at hour 0 was ∼10^7^, demonstrating cell viability at the time of inoculation. Strikingly, there was no significant decrease in culture CFU by hours 2 and 5. By hour 10 there was a slight decrease in CFU to ∼10^6^.. By 24 hours the culture CFU had significantly decreased to ∼10^3^.^5^, but surprisingly was not eliminated. From these data, we successfully determined the upper limit of *P. melaninogenica* growth to be between 5% and 8%, with no significant difference in maximum OD or cell viability at 5% compared to 0% O_2_. Second, the sustained presence of viable cells after 24 hours at 8% and 21% O_2_ demonstrates significant aerotolerance even in aerobic conditions.

**Fig 1.**
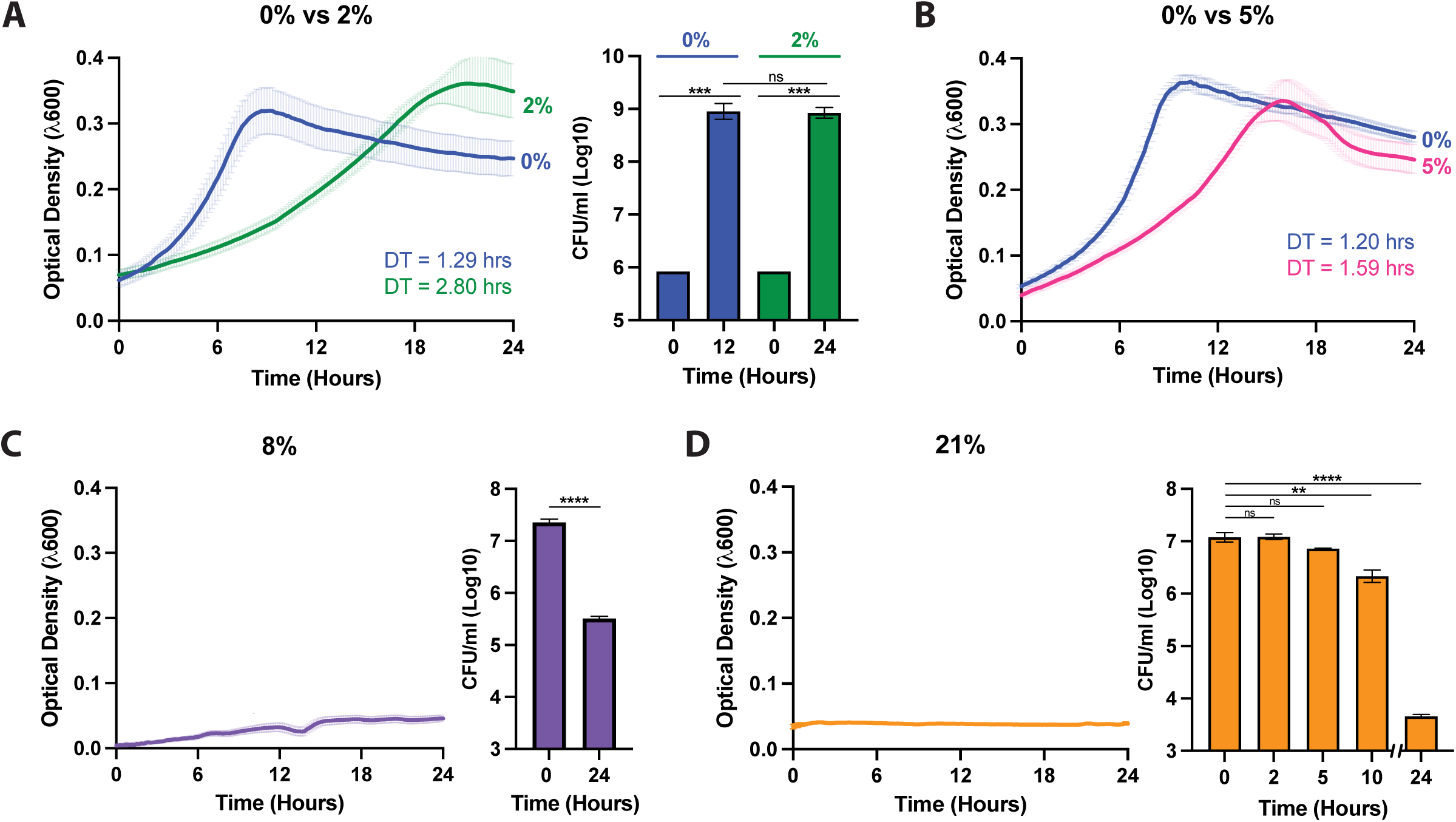
*Prevotella melaninogenica* grows in oxygen and remains tolerant when growth is inhibited at higher oxygen concentrations. A. *P. melaninogenica* simultaneously grown in 0% (blue) and 2% (green) over 24 hours. Optical density was measured at the wavelength 600. “DT” indicates the doubling time of each culture respectively. CFU at hour 0 vs time of the peak OD is shown for 0% (blue) and 2% (green) cultures with a bar graph. B. *P. melaninogenica* simultaneously grown in 0% (blue) and 5% (pink) over 24 hours. C. *P. melaninogenica* grown in 8% (purple) over 24 hours. CFU at hour 0 vs hour 24 is shown. D. *P. melaninogenica*Lorei mpsum grown in 21% (orange) over 24 hours. CFU plating was performed at hour 0,2,5,10 and.

Our next objective was to determine if *P. melaninogenica* consumes oxygen. This is necessary to better understand how *P. melaninogenica* is influencing its environment in the respiratory tract. Our approach was to determine the effect (if any) the presence of *P. melaninogenica* has on the surrounding oxygen concentration in liquid media. To do this, we measured the oxygen concentration and the cellular oxygen consumption rate (OCR) within the liquid cultures. The oxygen probe takes measurements of the O_2_ concentration and OCR every 30 seconds while moving up and down across 400um near the well floor. In this way, any gradient in O_2_ concentration correlating to a cell density gradient driven by cell settling will be quantified. The O_2_ concentration of liquid BHI media without cells was measured in parallel to confirm the desired O_2_ concentration fully diffuses into the media and that no O_2_ gradient is observed in the absence of cells. Similarly, the OCR of liquid BHI media without cells was measured in parallel to confirm that no OCR is observed in the absence of cells. The inoculum O_2_ concentration and cellular OCR was measured over 45 hours. When examining the O_2_ concentration in the liquid BHI cultures with *P. melaninogenica*, we find that the percent O_2_ varies significantly across well depths (Fig. 2A). This finding suggests that a lower oxygen level is correlated with a higher cell density as cell settling leads to higher cell density at the bottom of each well. When looking specifically at the representative 16-17 hour timepoint, we see that at the highest point in the probe’s 400um range (16hr:00min and 16hr:30min), the O_2_ concentration is ∼3%. However, at the bottom of the 400um range (16hr:15min and 16hr:45min), the O_2_ concentration is ∼2.4%, showing a striking 0.6% change in O_2_ over the span of 400um (Fig. 2B). In contrast, the media only control demonstrates no change in its 5% O_2_ concentration throughout the probe’s vertical scanning over the 16-17 hour timepoint (Fig. 2B). When examining the average O_2_ concentration of the media only control vs the media + *P. melaninogenica* culture over the entire 45 hour run, the overall change in oxygen level becomes extremely apparent, with the media only control demonstrating 5% O_2_ and the + *P. melaninogenica* culture demonstrating an average of 2.5% O_2_. Therefore, we found that the O_2_ decreases by∼2.5% in the presence of P. melaningenica (Fig. 2B). When examining the cellular oxygen consumption rate (OCR) in the liquid BHI cultures with *P. melaninogenica*, we find that the OCR is ∼50 (fmol/mm^2^/s) at 15 hours and that the OCR rises to ∼80 (fmol/mm^2^/s) by 45 hours (Fig. 2C). By contrast, the media only negative control shows no significant change in OCR from zero (Fig. 2C) When examining the average OCR of the media only control vs the media + *P. melaninogenica* culture over the entire 45 hour run, we again see that the OCR of the negative control is zero and the OCR of the *P. melaninogenica* culture is ∼50 (fmol/mm^2^/s). Based on these results, we determined that *P. melaninogenica* has a significant impact on its surrounding environment through the consumption of oxygen.

**Fig 2.**
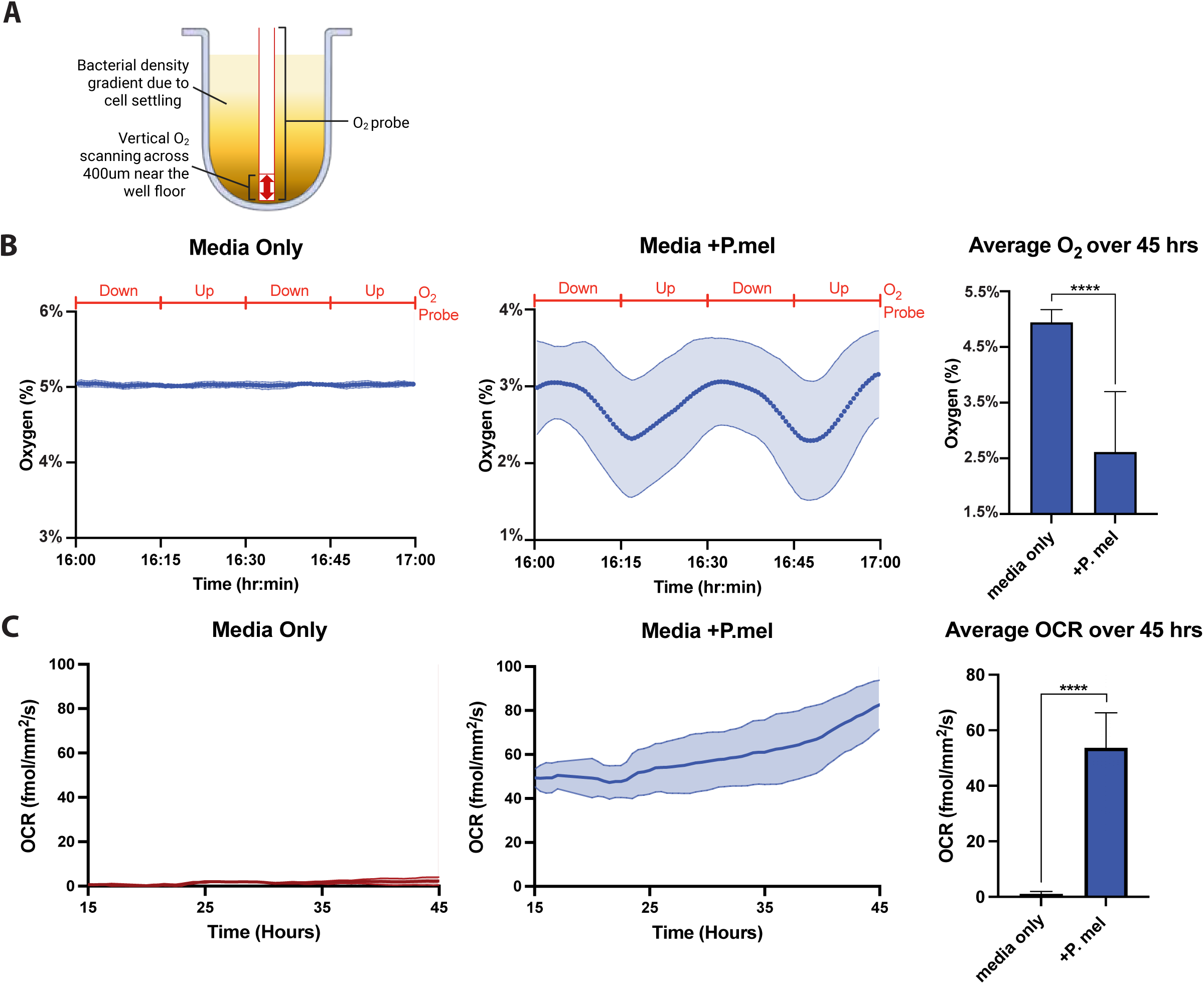
*Prevotella melaninogenica* consumes oxygen. A. Model of the Resipher oxygen probe used. A well with 100ul bacterial culture and the loaded O2 probe is pictured. The area of vertical scanning is represented with a red arrow. B. Oxygen measurements as the probe moves up and down across 400um over the span of one representative hour (hour 16 was chosen) in an atmosphere of 5% O2. Y axis is percent oxygen in the culture. X axis is time (hr:min). Red line above graphs represents the probe depth over the course of the hour. Left graph: O2 of BHI media – cells. Right graph: O2 of BHI media +*P. melaninogenica*. Bar graph depicts the average oxygen level of BHI media – cells and BHI media +*P. melaninogenica* over 45 hours. C. Oxygen Consumption Rate (OCR) quantification over time in an atmosphere of 5% O2. Y axis is the OCR (femtomole/square millimeter/seconds). X axis is time (hours). Left graph: OCR of BHI media – cells. Right graph: OCR of BHI media +*P. melaninogenica*. Bar graph depicts the average OCR level of BHI media – cells and BHI media +*P. melaninogenica* over 45 hours.

Given these striking observations, we then surveyed the genome to identify candidate genes that may be associated with oxidative stress defense in *P. melaninogenica,* based on the defense systems that have been worked out in the study of *E. coli*. This work enables a focused analysis, leveraging what is known about E. coli gene homologs to strengthen confidence in our findings. After the identification of E. coli oxidative stress genes, we then identified whether those genes were predicted to be present in the *P. melaninogenica* ATCC 25845 genome. This analysis resulted in the identification of 16 genes (Table 1). All genes are protein coding, therefore along with the ENSEMBL gene IDs, the protein symbols and names are listed. The 16 genes and their associated protein symbols are: *nqrA* (NqrA), *nqrB* (NqrB), *nqrC* (NqrC), *nqrD* (NqrD), *nqrE* (NqrE), *nqrF* (NqrF), *appC* (CydA), *cydB* (CydB), *ahpF* (AhpF), *ahpC* (AhpC), *rbr* (Rbr), *bcp*(Bcp), *dfx* (Dfx), *ftnA* (FtnA), *rd* (Rd), and *oxyR* (OxyR). Therefore, we developed a list of 16 genes predicted to be involved in the oxidative stress defense system in *P. melaninogenica*.

**Table 1.**
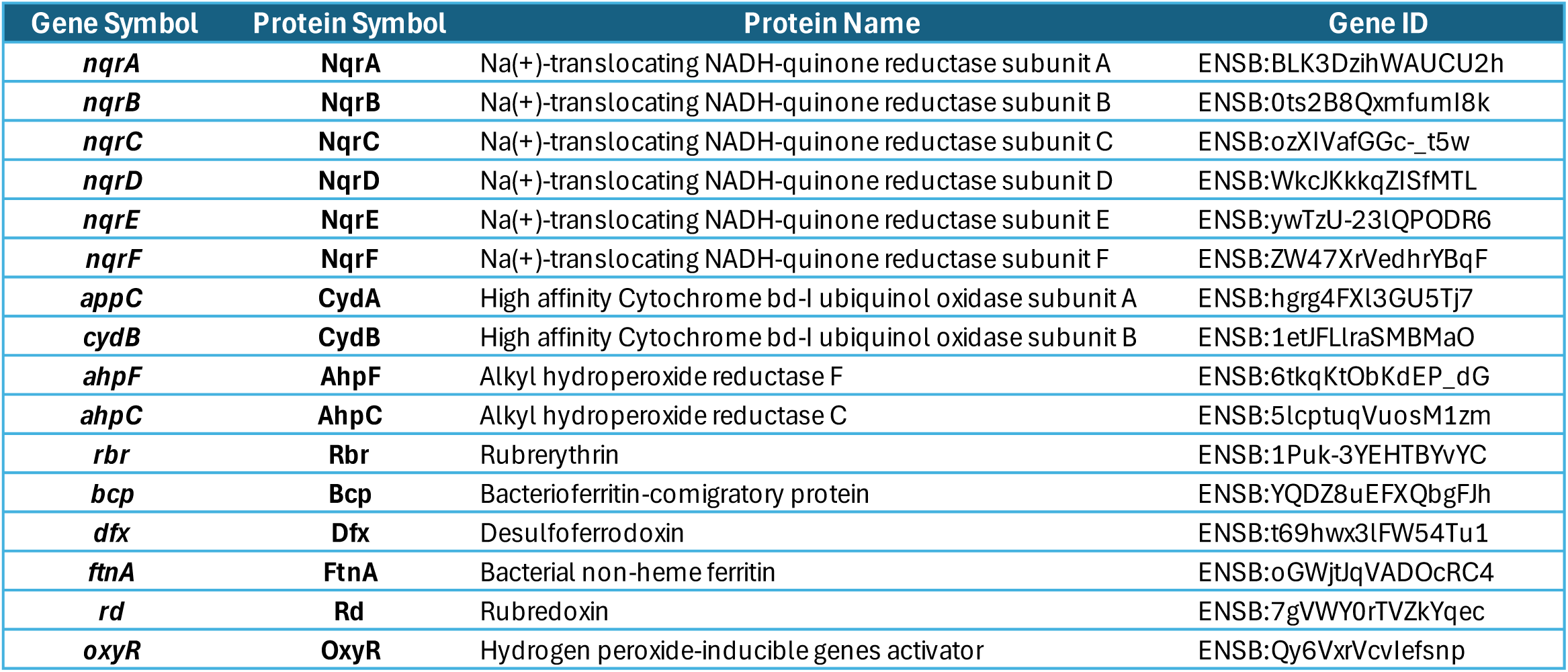
Putative oxidative stress defense genes in *P. melaninogenica*. Gene symbol, protein symbol, protein name and ENSEMBL gene IDs are listed.

Our next objective was to determine if the genes predicted to be associated with oxidative stress defense are significantly upregulated under low oxygen conditions compared to anaerobic conditions. We performed total RNA sequencing to quantify the transcriptomic profile of *P. melaninogenica* grown within 2% vs. 0% O_2_ environments. The inoculating culture of *P. melaninogenica* was initially grown in Brain Heart Infusion (BHI) broth at 37°C under strictly anaerobic conditions (0% O_2_). Inoculums derived from this culture were then cultured in BHI broth at 37°C in 2%O_2_ and 0% O_2_ simultaneously. Samples were all collected during early exponential phase with ODs of ∼0.35. Analysis focusing specifically on our 16 genes of interest revealed that *all* candidates were significantly upregulated in the 2% O_2_ condition compared to their respective anaerobic (0% O_2_) expression levels. Genes *ahpF and ahpC were* the two most significantly upregulated genes by a substantial margin. Following ahpF and ahpC, listed in descending order of expression, was: *appC*, *cydB, rd*, *dfx*, *ftnA*, *bcp*, *nqrA*, *rbr*, *oxyR*, *nqrD*, *nqrC*, *nqrF*, *nqrB*, and *nqrE* (Fig 3 and Sup. Table 1). These results demonstrate that the genes we predicted to be associated with oxygen defense and tolerance are in fact significantly upregulated in the low oxygen 2% condition.

**Fig 3.**
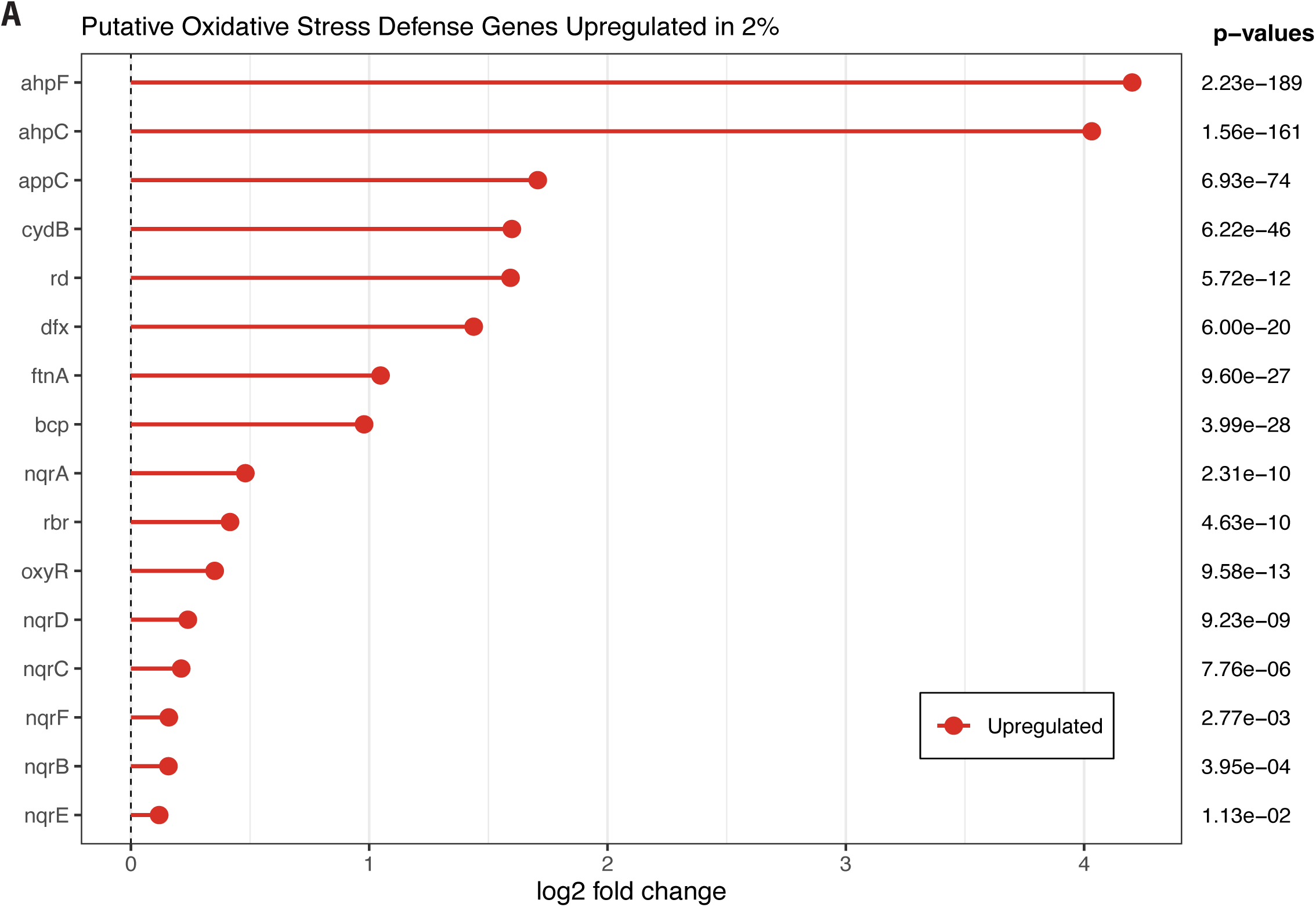
All putative oxidative stress defense genes are significantly upregulated in 2% oxygen vs. 0% oxygen. Y axis is labeled with gene symbols. X axis is log 2 fold change from gene expression in 0% O2. Adjusted p values of each respective gene are listed.

Our next objective was to identify if there were any groups of significantly upregulated genes with functions outside oxidative stress defense. In the transcriptomic dataset described above, We found 17 genes associated with DNA protection and repair in E. coli were significantly upregulated in *P. melaninogenica* growing in 2% O_2_ when compared to the basal anaerobic expression levels (Fig 4A). The log2 fold change of each upregulated gene is listed in Supplemental Table 2. Additional information for each of the 17 genes is shown in a table (Fig 4B). All 17 genes are protein-coding, therefore the protein symbols and names are shown in addition to the gene ENSEMBL IDs. These results demonstrate that there are many genes associated with DNA protection and repair that are significantly upregulated in 2% oxygen.

**Fig 4.**
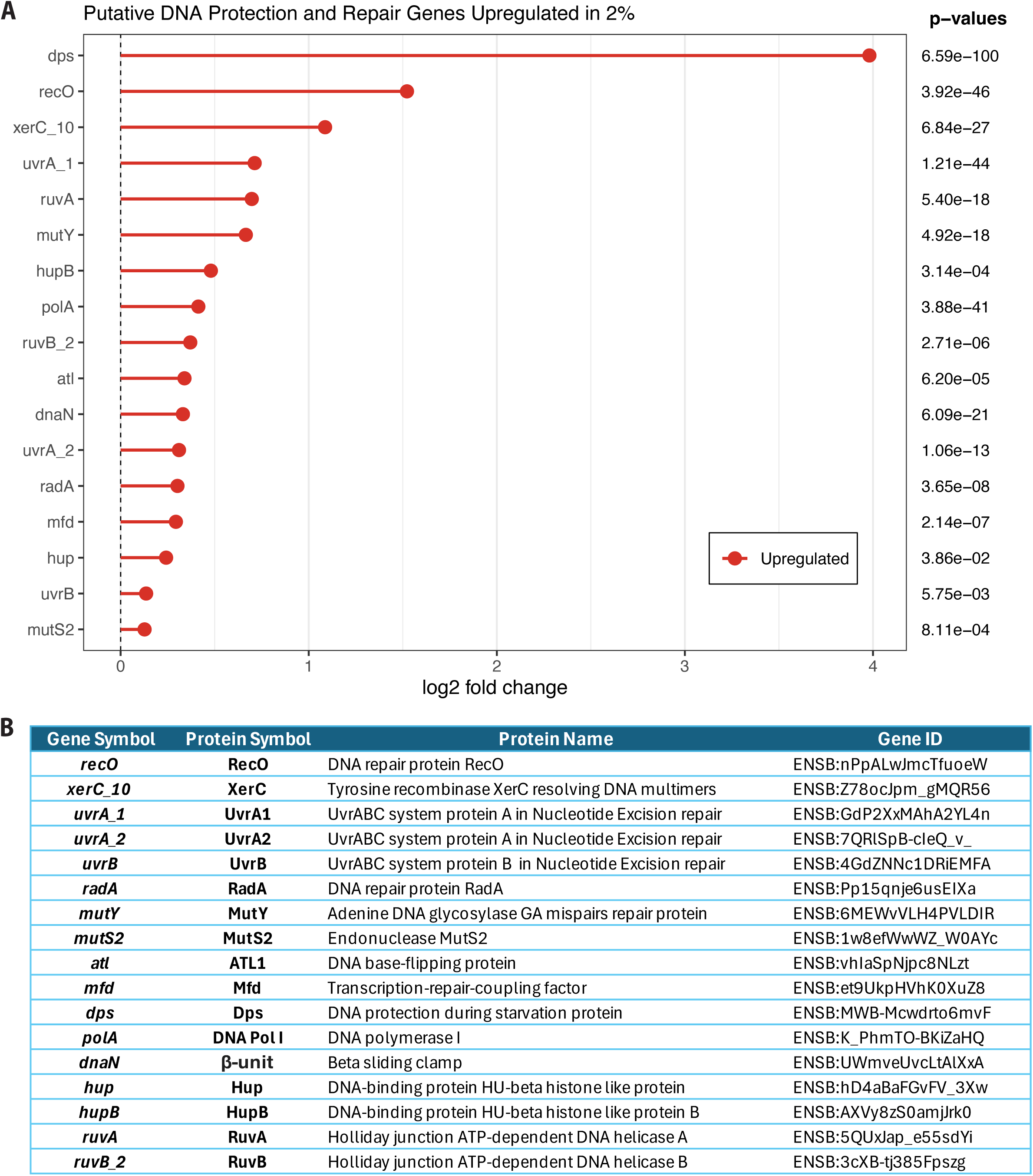
Putative DNA Integrity and Repair genes are significantly upregulated in 2% oxygen vs. 0% oxygen. A. Upregulated genes with gene labels on the Y axis and log 2 fold change on the x axis. Adjusted p values of each respective gene are listed. B. Putative integrity and repair gene table based on the transcriptomic results. Gene symbol, protein symbol, protein name and ENSEML ID are shown.

Finally, we sought to identify protein homologs across the *Prevotella* genus and formerly classified Prevotella species to the identified putative oxidative stress defense proteins in *P. melaninogenica.* Due to the association of the *Prevotella* genus with the highly oxic respiratory tract and the association of reclassified *Prevotella* species with the lower-oxygen vagina and gut, we sought to examine the potential association of oxidative stress defense machinery and habitat oxygen levels. To determine homology, we performed a BLASTp analysis of each identified putative oxidative stress defense protein in *P. melaninogenica* against genomes of the 21 *Prevotella* species and 13 reclassified species commonly found in the gut and vagina. Protein hits within each species with >60% amino acid coverage and <0.05 E value were considered homologs and included in our analysis. The degree of homology was determined via amino acid similarity scores. Protein homologs were depicted with a heatmap, where the amino acid similarity is represented by color (100% similarity = orange; 0% similarity = ivory). The bacteria were then grouped into a dendogram based on the homology results. Across all 21 *Prevotella* species, 11 of the 16 analyzed proteins demonstrated amino acid similarity to those in *P. melaninogenica*: NqrB, NqrC, NqrD, NqrE, NqrF, CydB, AhF, AhpC, Bcp, Dfx, and Rd (Fig 6). The average amino acid similarity for each protein across the genus was between 81-97%. However, several proteins exhibited lineage-specific absence. NqrA homologs with >82% AA similarity were conserved in 20 species but were absent in *P. falsenii* (0% AA similarity). CydA homologs (>87% AA similarity) were identified in all species except *P. dentasini* and *P. falseniil* (0% AA similarity). Rbr homologs (>84% AA similarity) were absent in *P. dentasini* and *P. micans* (0% AA similarity). FtnA homologs (>78% AA similarity) were not detected in *P. micans* and *P. nigrescens* (0% AA similarity). Finally, OxyR homologs (>81% AA similarity) were present in all species with the exceptions of *P. brunnea* and *P. dentasini* (0% AA similarity). Hierarchical clustering of *Prevotella* and reclassified *Prevotella* species based on protein similarity heatmaps revealed six distinct groupings (Fig 6). These groupings demonstrate a progressive increase in AA similarity relative to *P. melaninogenica*. The first group to diverge consisted of *Segatella bryantii*, *S. albensis*, *Xylanibacter rodentium*, *X. brevis*, and *X. ruminicola*; a distinguishing characteristic of this group is the near complete absence of Dfx and FtnA homologs. *P. dentasini* and *P. falsenii* formed the second group, while *Leyella stercorea*, *P. micans*, and *P. nigrescens* constituted the third. The fourth group comprised *S. copri*, *P. brunnea*, and *S. hominis*. A large intermediate group separated prior to the *P. melaninogenica* group, containing *Hallela bergensis*, *Hoylesella buccalis*, *Hoylesella timonensis*, *Hallela colorans*, *Hallela mizrahii*, *P. amnii*, *P. bivia*, *P. corporis*, *P. disiens*, *P. pallens*, *P. aurantiaca*, and *P. intermedia*. The final group, inclusive of *P. melaninogenica*, contained nine species: *P. veroralis*, *P. vespertina*, *P. histicola*, *P. denticola*, *P. multiformis*, *P. fusca*, *P. jejuni*, *P. scopos*, and *P. melaninogenica*. This group exhibited a mean AA similarity of 96.3%. From these data, we successfully determined protein homology to the identified putative oxidative stress defense proteins in *P. melaninogenica* across the Prevotella genus and formerly classified Prevotella species.

**Fig 5.**
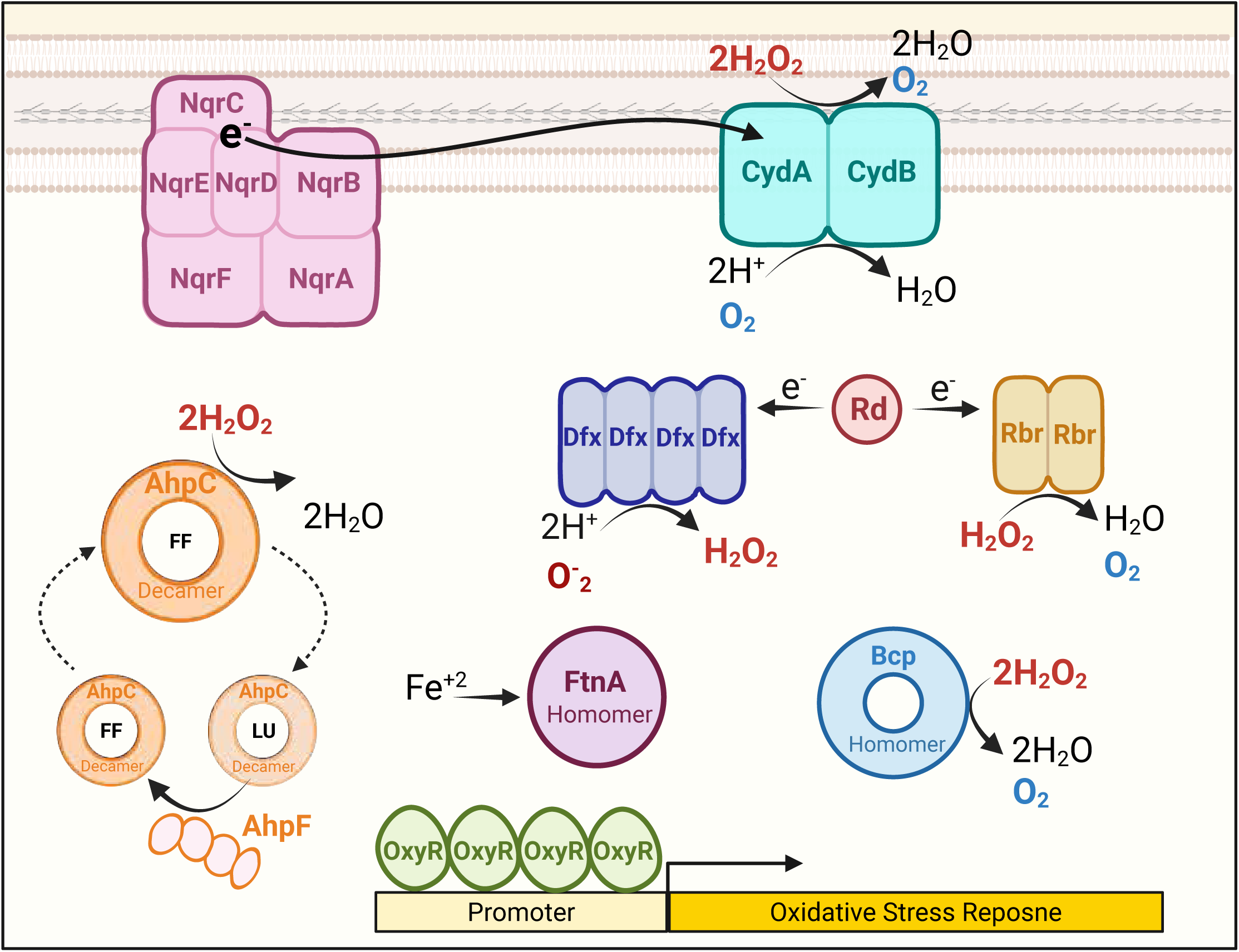
The putative oxygen defense mechanisms upregulated in 2% oxygen to confer tolerance. The membrane associated Na+-NQR complex is shown in pink, with the six subunits NqrABCDEF labeled in dark pink. The membrane associated High-affinity Cytochrome bd-I (Cyd bd I) is shown in aqua, with its two subunits CydAB labeled in dark aqua. Rubredoxin (Rd) is a red circle, depicting a monomer. Desulfoferrodoxin (Dfx) is a dark blue tetramer complex. Rubrerythrin (Rbr) is a yellow dimer. AhpC is in orange, representing a ring-like decamer. AhpC’s two active site conformations, (fully folded and locally folded) are represented with “FF” and LU”. AhpF is in orange, representing a homodimer. Bacterio-ferritin-comigatory protein (Bcp) is a blue ring-like homomer (a complex with many identical subunits). Bacterial non-heme ferritin (FtnA) is a maroon circle which represents another homomer. Finally, the active OxyR tetramer is shown in green on the promoter of the oxidative stress gene operon. Electron flow is depicted with a black arrows. Reactive oxygen species (ROS) are depicted in red. Oxygen is depicted in blue.

**Fig 6.**
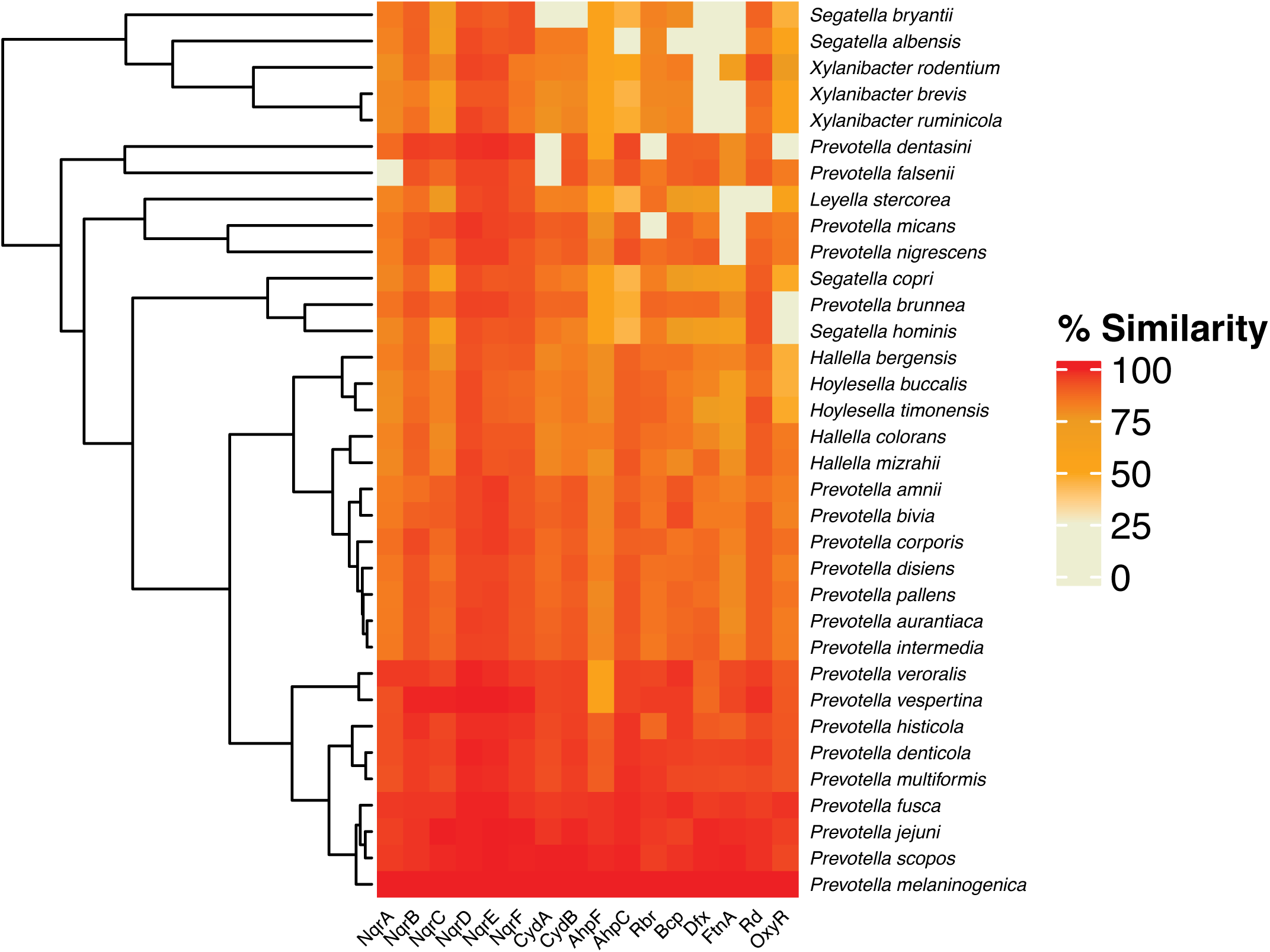
Homologs of the identified putative oxidative stress defense proteins upregulated in *P. melaninogenica* are nearly ubiquitous across the *Prevotella* genus but less prevalent in the reclassified *Prevotella* species. The y axis denotes the 21 *Prevotella* species (primarily associated with the respiratory tract), and 13 reclassified *Prevotella* species (associated with the gut and vagina). The x axis denotes proteins encoded by the 16 upregulated putative oxidative stress defense genes in *P. melaninogenica*. The top BLASTp hit with >60% coverage and <.05 E value within each bacteria for each protein is shown. The percent amino acid similarity of each candidate homolog is depicted with a color gradient (red = 100%; cream = 0%). Bacteria are clustered based on protein homology.

## METHODS

### P. melaninogenica reconstitution and culture conditions

*Prevotella melaninogenica* 25845 was purchased from ATCC as a frozen culture. Bacteria was grown up in Brain Heart Infusion (BHI) broth (Anaerobe Systems) broth under anaerobic conditions in an AS-500 anaerobic chamber (Anaerobe Systems). Gram stain and Sanger sequencing was performed to confirm ID.

### *P. melaninogenica* growth and viability assay

For each respective run, the inoculating culture of *P. melaninogenica* was initially grown in Brain Heart Infusion (BHI) broth at 37°C under strictly anaerobic conditions (0% O_2_). All inoculums derived from this culture were then cultured in BHI broth at 37°C in either the anaerobic chamber for 0% O_2_, the hypoxic chamber (Coy) for 2%, 5% and 8% O_2_ respectively, or ambient air for 21% O_2_. Upon detection of *P. melaninogenica* growth in 2% O_2_ and 5% O_2_ respectively, the assays were repeated with the addition of parallel 0% O_2_ cultures inoculated from the same parent culture for direct comparison of growth rate, maximum optical density (OD), and cell viability over time in 0% vs 2% O_2_ and 0% vs 5% O_2_. Growth kinetics were quantified by measuring optical density (OD) over time at wavelength 600 using AltoMicroplate Readers (Cerillo). For parallel anaerobic and aerobic assays, identical 96-well plates were read by a microplate reader in the anaerobic chamber and hypoxic chamber (at either 2% or 5% O2) respectively. Cell viability was quantified via colony-forming units (CFU) following anaerobic plating. All cultures were plated at hour 0 to establish baseline inoculum viability. For each respective run, culture was plated anaerobically at hour 0 to quantify the number of viable cells in the inoculum via colony-forming units (CFU). Subsequent anaerobic plating was performed at the time of maximum OD or, in the absence of observed growth, at 24 hours. For cultures exposed to 21% O_2_, a death curve was established by plating at 0, 2, 5, 10 and 24 hours. Bacterial identity was confirmed using Gram stain and PCR.

### Quantifying Significance

An un-paired t test with Welch’s correction was performed to determine p values. P values of <.05 were considered significant. P value range is indicated with 1-4 stars.

### Measuring environmental oxygen and cellular oxygen consumption rate

The Resipher system (Lucid Scientific) was used to quantify the soluble oxygen and cellular oxygen consumption rate (OCR) over time and at varying well depths. The Resipher system consists of a 96-well plate containing inoculums, a sensing lid with probes, the processing *device*, and the base station *hub.* Each probe is coated in a material which, when optically excited, produces fluorescence and light directly proportional to the level of oxygen in solution. In this way, the O_2_ concentration is quantified. The OCR is calculated based on the fmol O_2_ /mm^2^/second. Measurements are taken every 30 seconds while the probe moves up and down across 400um near the well floor. One full cycle of vertical scanning takes 15 minutes.

For each assay, the inoculating culture of *P. melaninogenica* was initially grown in Brain Heart Infusion (BHI) broth at 37°C under strictly anaerobic conditions (0% O_2_). 100ul BHI inoculums derived from this culture and 100ul BHI negative controls were loaded into the center wells of a Cornell round bottom 96-well plate under strictly anaerobic conditions (0% O_2_). Outer wells were loaded with PBS to mitigate evaporation. The 96-well plate was then transferred to the hypoxic chamber via an anaerobic transfer container. We let the plate sit for 1 hour at 37°C to allow for cell settling, acclimation to temperature and acclimation to the set atmospheric oxygen (2%, 5% or 8% respectively). One hour prior to this incubation, the Resipher device and sensing lid were placed in the hypoxic chamber to acclimate for 2 total hours prior to being loaded onto the 96-well plate. After the lid and device were loaded onto the plate, the assay was run for 45 hours.

### Sample preparation and RNA sequencing

The inoculating culture of *P. melaninogenica* was initially cultured in Brain Heart Infusion (BHI) broth at 37°C under strictly anaerobic conditions (0% O_2_). Five biological replicates (n=5) were established in BHI broth with a standardized starting optical density (OD_600_) of 0.05. Two identical aliquots were made from each biological replicate. These two sets of 5 inoculums were cultured simultaneously at 37°C under either 2% O_2_and 0% O_2_ conditions. Cells were harvested during exponential phase, with an average sample OD of ∼0.36. Cultures were stored according to the Qiagen RNAprotect protocol until further processing. Total RNA was isolated using Qiagen RNeasy Mini Kit. Quality control analysis confirmed that all samples met library standards, with DV200 values between 71-93% and ≥100ng total RNA. Total RNA library prep with ribonucleotide-depletion was performed. Short-read sequencing was performed using the Aviti24 high-output kit. Libraries were sequences to a depth of ∼20 million reads/sample.

### RNA sequencing and Analysis

Raw paired-end RNA-seq reads were processed using fastp (version 1.0.1) for adapter detection and quality trimming. The software automatically identified paired-end adapter sequences (--detect_adapter_for_pe) and performed 5′ and 3′ sliding-window trimming to remove low-quality bases (--cut_front, --cut_tail, --cut_window_size 4, --cut_mean_quality 20). Poly-G homopolymers were trimmed (--trim_poly_g), and reads shorter than 50 bp post-trimming were discarded.A transcriptome index for *Prevotella melaninogenica* ATCC 25845 (assembly GCA_000144405) was constructed using Salmon (version 1.10.3) with a k-mer size of 31. Trimmed reads were quantified using Salmon. For each sample, paired fastp-processed reads were supplied to salmon quant with selective alignment–based validation (--validateMappings), and GC-bias correction (--gcBias). Estimated counts from Salon were combined into a gene count matrix.Gene-level count matrices were imported into R and analyzed using DESeq2 (version 1.46.0). Low-count genes were filtered according to DESeq2 defaults, size factors were estimated using the median-ratio method, and dispersion parameters were fit using DESeq2’s standard workflow^17,18,19^.

### Protein Homology Analysis

BLASTp analysis of each identified putative oxidative stress defense protein in *P. melaninogenica* against genomes of the 21 Prevotella species and 13 reclassified species commonly found in the gut and vagina was performed. Protein hits within each species with >60% amino acid coverage and <0.05 E value were considered homologs and included in our analysis. The degree of homology was determined via amino acid similarity scores. Homologs were depicted with a heatmap, where the amino acid similarity is represented by color (100% similarity = Orange; 0% similarity = Ivory). The bacteria were then grouped into a dendogram based on the homology results.

## DISCUSSION

This study resolves a biological paradox by demonstrating that *Prevotella melaninogenica*, traditionally classified as a strict obligate anaerobe, possesses the capacity for robust growth and survival in oxygenated environments. Our results indicate that while the upper limit for active growth is between 5% and 8% oxygen, the organism exhibits significant aerotolerance even at atmospheric levels (21% O_2_), maintaining viable subpopulations for at least 24 hours. We propose that this survival is underpinned by the organism’s active modification of its local environment through oxygen consumption and a sophisticated, oxygen-induced transcriptional response. Specifically, we observed the significant upregulation of 16 putative oxidative stress defense genes—such as the Na^+^-NQR (*nqrABCDEF*), the high-affinity cytochrome *bd I* oxidase (*cydAB*), and the superoxide reductase (SOR) Desulfoferrodoxin (*dfx*)—alongside 17 genes dedicated to DNA protection and repair, most notably *dps*. Furthermore, protein homology analysis reveals that these oxidative defense mechanisms are highly conserved across the *Prevotella* genus, particularly among species inhabiting the human respiratory tract. Collectively, these findings provide a necessary correction to the century-old classification of *P. melaninogenica* and provides evidence that its persistence in the oxygenated human airway is likely driven by a specialized molecular toolkit for oxygen defense and DNA maintenance.

Our data demonstrate that *P. melaninogenica* proliferates robustly in 2% and 5% oxygen, achieving a maximum optical density and cell viability comparable to anaerobic controls with only a marginal reduction in replication rate. This discovery challenges a century of classification, beginning with Oliver and Wherry (1921), who designated the organism as an obligate anaerobe based primarily on its inability to grow in ambient air. The classification of strict anaerobe has been upheld by a reliance on binary experimental frameworks—comparing only 0% to 21% O_2_—while leaving intermediate concentrations largely unexplored^9^. Early attempts to examine modular oxygen levels, such as those by Rosenblatt (1975) and Loesche (1969), were hindered by unmeasured oxygen concentrations, qualitative metrics, or the use of non-standardized clinical strains, making their results unreliable. Furthermore, previous methodologies often focused on transient oxygen exposure or colony morphology on solid media, which does not capture the comprehensive growth kinetics provided by our simultaneous and continuous liquid culture monitoring. Therefore, this study establishes for the first time that *P. melaninogenica* growth at 2% and 5% oxygen is sustained and robust from the point of inoculation. These findings suggest that the physiological growth threshold of this species is significantly higher than previously recognized, necessitating a revision of its classification as a strict anaerobe.

Our results demonstrate that even when active proliferation is arrested at elevated oxygen concentrations, *P. melaninogenica* maintains sustained aerotolerance for at least 24 hours. Notably, in ambient air (21% O_2_), no significant decrease in viability was observed during the first 10 hours of exposure, with a persistent subpopulation surviving through the full 24-hour period. While early observations by Rosenblatt (1975) suggested that certain clinical isolates might tolerate ambient air for up to two days, the use of non-standardized strains and qualitative plate-based metrics limited the broader applicability and reproducibility of those findings. Our quantitative data are more robustly supported by the physiological model of the closely related *Bacteroides fragilis*; although its growth threshold is significantly lower than that of *P. melaninogenica* (0% – 0.14% O_2_) ^10^, it exhibits a comparable capacity to withstand 24–72 hours of atmospheric exposure^12,13^. Interestingly, in *B. fragilis*, growth arrest in oxygenated conditions is a regulated process involving the *oxe* gene, suggesting that the cessation of division at higher concentrations may not be a failure of oxidative stress defense, but rather a strategic entry into a dormant state to prioritize survival. In our study, colonies from the anaerobic plating performed after a *P. melaninogenica* liquid culture was exposed to 21% O_2_ for 24 hours were significantly smaller than those plated at the same time point from a liquid culture grown anaerobically. This observation supports our hypothesis that P. melaninogenica enters cell dormancy as an adaptive strategy, as it would likely take a longer time to exit this state and restart replication after being placed back into an environment conducive for growth. In the context of the human lung—where oxygen tension varies dramatically from approximately 9.9% O_2_ in healthy airways to as low as 1% within the thickened mucus of cystic fibrosis patients—this physiological flexibility is critical^8^. The capacity of *P. melaninogenica* to grow robustly at 5% while remaining resilient at 21% establishes it as an organism highly adapted to the spatial and pathological heterogeneity of the respiratory tract.

Beyond growth and tolerance, our results demonstrate that *Prevotella melaninogenica* actively modifies its local environment through significant oxygen consumption, with measured rates between 50–80 fmol O₂/mm²/s in 2% oxygen. The capability to consume oxygen has not been studied in any *Prevotella* species, therefore these findings fill a gap in the literature regarding the metabolic capabilities of these commensals. While oxygen consumption has been previously characterized in *Bacteroides fragilis*—which exhibits a consumption rate of 50 nmol/min (9 nmol/min/mg dry cells) in 0.1% O_2_^10^ —our findings extend this metabolic trait to *P. melaninogenica* under significantly higher oxidative pressures. The ability to consume oxygen likely serves as an evolutionary adaptation to enable colonizaton of niches where oxygen concentrations would otherwise overwhelm the bacterium’s oxidative stress defense systems. In the context of the highly variable oxygen gradients of the human lungs, this strategy would provide a distinct advantage, likely explaining the predominance of *P. melaninogenica* in the lower airway. Furthermore, it is likely that the localized depletion of oxygen by *P. melaninogenica* plays a role in the broader microbial landscape. Seeing as *P. melaninogenica* is frequently reported with other taxa like *Veillonella and Fusobacterium spp.*, these species may rely on the protective, low-oxygen microenvironments created by *P. melaninogenica*. Our data provide novel insight into the oxygen-scavenging capabilities of *P. melaninogenica* at 2% O_2_, identifying a mechanism of environmental niche modification. These findings have implications for respiratory microbial ecology, suggesting that *P. melaninogenica* may act as a keystone species that facilitates the survival of complex polymicrobial communities through the modulation of local redox conditions.

Our results show that two protein complexes associated with aerobic respiration in low-oxygen environments, NADH:quinone oxidoreductase (Na^+^-NQR) and the high-affinity terminal oxidase Cytochrome bd-I CydAB, are significantly upregulated in *P. melaninogenica* under 2% O_2_. Historically, members of the *Prevotella* genus were considered limited to fermentation and anaerobic respiration due to their classification as anaerobes^20,21^. However, genomic sequencing has revealed that several supposedly anaerobic bacteria, such as *B. fragilis*, possess oxygen-directed respiratory systems^22^. These low-oxygen respiratory systems comprise an NADH dehydrogenase (NADH2 and/or Na^+^-NQR), a quinol electron carrier, and a high-affinity terminal oxidase (CydAB), which catalyzes the reduction of oxygen to water. In contrast, anaerobic respiration utilizes fumarate reductase as the terminal oxidase to catalyze the reduction of fumarate to succinate in the absence of oxygen. In *B. fragilis*, Na^+^-NQR and NADH2 contribute approximately 77% and 23%, respectively, to the quinol pool under low-oxygen conditions^10,15^. Protein expression analysis of fumarate reductase and CydAB in *B. fragilis* under anaerobic and low-oxygen conditions indicates that fumarate reductase expression is higher at 0% O_2_ but remains significantly upregulated in low oxygen. Conversely, CydAB is highly expressed under low-oxygen conditions with negligible expression under anaerobic conditions. Both terminal oxidases utilize the electron carrier menaquinone from the quinone pool^10,15^. To validate these associations, a fumarate reductase knockout (Δfrd) was grown in anaerobic and low-oxygen conditions. The lack of growth in Δfrd under anaerobic conditions suggests that CydAB is ineffective for energy production without oxygen and that fermentation does not supplement anaerobic respiration. However, significant growth of Δfrd in low oxygen indicates that CydAB activity drives energy production through aerobic respiration at low oxygen levels^10,15^. The *P. melaninogenica* genome is predicted to encode Na^+^-NQR, NADH2, menaquinone, fumarate reductase, CydAB, and ATP synthase. Our transcriptomic data show that while NADH2, menaquinone, and fumarate reductase are transcribed under anaerobic conditions, only Na^+^-NQR and NADH2 are significantly more expressed at 2% oxygen (Fig 5). These findings align with the expression patterns observed in *B. fragilis* outlined above. Therefore, we propose that under 2% O_2_, *P. melaninogenica* energetically benefits from low levels of oxygen through a type of aerobic respiration.

In addition to energetic concerns, microbes growing in oxygen must also have a strategy to mediate oxidative stress. Our genomic and transcriptomic analysis of *P. melaninogenica* also shows that superoxide dismutase (SOD) is not predicted to be present, yet a superoxide reductase (SOR) identified as desulfoferrodoxin (*dfx*) is present and significantly upregulated in 2% O_2_. SOD is a well-characterized superoxide scavenger in many bacteria, including *B. fragilis* and *E. coli* ^23^. The absence of SOD in the *P. melaninogenica* genome is significant given the enzyme’s importance in ROS detoxification. However, an alternative superoxide scavenger, SOR—previously identified in *Desulfoarculus baarsii*^24^, and *Desulfovibrio vulgaris*^25^—is present in *P. melaninogenica* and upregulated under 2% O_2_. This finding is particularly notable as SORs have not been identified in the closely related *Bacteroides* genus. Further genomic analysis of the SOR *dfx* reveals that it is predicted to be present in other *Prevotella* species such as *P. histicola* and *P. intermedia*, suggesting this may represent a distinct feature of the *Prevotella* genus. SOD and SOR are redox-active metalloenzymes which have evolved independently and utilize different catalytic mechanisms. While SOD catalyzes the disproportionation of O_2_^•–^ into O_2_ and H_2_ O_2_, SOR catalyzes the single-electron reduction of O_2_^•–^ to H_2_O, using two protons and the electron donor rubredoxin (also identified in this study to be upregulated in 2% O_2_)^23,25^. SODs consume two superoxide anions per cycle and SORs consume one, however the SOR reaction involves the simultaneous consumption of NAD(P)H. This process diminishes the overall negative redox status of the cell, which ultimately reduces intracellular superoxide production^26^. The biological implication of SOD versus SOR in bacteria has yet to be elucidated, however, it has been proposed that the absence of O_2_ production may be the primary advantage of SOR over SOD, as reducing the quantity of free oxygen within the cell likely prevents the further generation of ROS^23^. Our data showing the significant upregulation of SOR under aerobic conditions supports its role in superoxide detoxification (Fig 5). These results suggest that *P. melaninogenica* may utilize unique elements within its ROS detoxification machinery that distinguish it (and its oxygen tolerance) from the related *Bacteroides* genus.

Beyond its role in energy generation, cytochrome bd-I (CydAB) has been proposed to enable a number of other vitally important physiological functions. CydAB was found to have peroxidase activity in E. coli^27^, enhance tolerance to nitrosative stress in E. coli^28^, and most notably, drive the consumption of oxygen in *B. fragilis*^10^. Given the robust oxygen consumption by *P. melaninogenica* observed in our study, experimental validation of CydAB’s role in this mechanism is grounds for future study. Consequently, we propose that the significant upregulation of CydAB in P. melaninogenica is likely driven by both its energetic benefits and potential contribution to oxidative stress defense^29^.

Our results show that AhpC and AhpF are the most significantly upregulated genes in *P.melaninogenica* under 2% oxygen, with differential expression levels markedly higher than any other significantly upregulated genes. These genes encode peroxiredoxins (Prxs), a peroxidase family essential for antioxidant protection in diverse bacteria^30^. In *E. coli*, peroxiredoxins are among the ten most abundant proteins, supporting our transcriptomic findings, and suggesting that these proteins serve as primary mediators of the cellular oxidative stress response^31^. Mechanistically, AhpC and AhpF function as a synergetic unit to catalyze the NADH-dependent reduction of H_2_O_2_ to H_2_O in *E. coli*^30^. AhpC serves as the peroxidase that directly reduces H_2_O_2_, while AhpF acts as the reductase required to maintain AhpC in its active, fully folded conformation^30^. The exceptional fold-change observed in our data, combined with the established ROS-scavenging roles of these enzymes in *E. coli*, indicates that this conserved system is a critical functional priority for *P. melaninogenica*. These results suggest that AhpC/F activity is central to maintaining redox homeostasis within the oxygen-variable environment of the human respiratory tract (Fig 5).

In addition to putative oxidative stress machinery, we identified many putative DNA repair genes upregulated in 2% O_2_, indicating a previously unknown damage repair response in addition to O_2_ defense in *P. melaninogenica*. In *E. coli,* strains that are deficient in both recombination and excision repair strategies are fully viable only in anaerobic media, thus DNA repair was found to be an essential component of the oxidative stress response for any growth in oxygen^32^. The DNA repair genes upregulated in *P. melaninogenica* are predicted to fall into the following functional groups: DNA binding protein (*dps*), nucleotide excision repair (*uvrA_1, uvrA_2,* and *uvrB*), holiday junction helicase (*ruvA* and *ruvB_2),* the primary DNA repair response (*recO*), GA mismatch repair (*mutY*), DNA base flipping (*atl*), histone like (*hupB* and *hup*), DNA polymerase (*polA*), mutation frequency decline (*mfd*), recombination processing (*radA*), DNA multimer resolution (*xerC_10*), endonuclease (*mutS2*), and beta sliding clamp (*dnaN*). In *E. coli*, DNA repair is regulated by the SOS system^32^. However, this regulator was not found to be upregulated in *P. melaninogenica*. No other proteins previously characterized as DNA repair regulators were found to be upregulated in *P. melaninogenica*, however the genome of *P. melaninogenica* is not well characterized, resulting in many genes remaining unannotated. It is likely that the regulator of DNA repair in *P. melaninogenica* is a novel protein in need of characterization. This is therefore grounds for future investigation.

Finally, our protein similarity analysis revealed that the oxidative stress defense proteins upregulated in *P. melaninogenica* are nearly ubiquitous across the current members of the *Prevotella* genus, yet less prevalent in recently reclassified species. Given the evidence for robust oxidative defense machinery identified in our data, we investigated whether this repertoire was unique to *P. melaninogenica* or a conserved trait within the newly reorganized genus. Due to the high degree of interspecies diversity, many bacteria formerly within *Prevotella* were recently reclassified^33^. The refined *Prevotella* genus now comprises 21 species primarily localized to the human respiratory tract, whereas many reclassified taxa occupy niches with lower oxygen availability, such as the gut and vaginal microbiomes. The finding that reclassified species possess homologs of lower similarity compared to true *Prevotella* suggests a correlation between environmental oxygen levels and the specific repertoire of oxidative defense machinery. Furthermore, the presence of protein homologs with high amino acid similarity across the genus indicates that the capacity for oxygen growth and tolerance is likely a shared trait rather than one unique to *P. melaninogenica*. These results necessitate a re-evaluation of the physiological classification of the *Prevotella* genus as a whole. This study represents a critical first step in elucidating the adaptive physiology and microbial dynamics of *Prevotella melaninogenica* within the oxygen-variable landscape of the human lung microbiome.

## Data Availability Statement

Sequences related to this submission will be made available as an NCBI BioProject. BioProject accession number PRJNA1400211

